# Heterogeneous Kinetics of Nanobubble Ultrasound Contrast Agent and Angiogenic Signaling in Head and Neck Cancer

**DOI:** 10.1101/2024.09.22.614362

**Authors:** Benjamin Van Court, Mark Ciccaglione, Brooke Neupert, Michael W. Knitz, Sean P. Maroney, Diemmy Nguyen, Khalid N.M. Abdelazeem, Agata A. Exner, Anthony J. Saviola, Richard K.P. Benninger, Sana D. Karam

## Abstract

Recently developed *nanobubble* ultrasound contrast agents are a promising tool for imaging and drug delivery in tumors. To better understand their unusual kinetics, we implemented a novel pixel clustering analysis, which provides unique information by accounting for spatial heterogeneity. By combining ultrasound results with proteomics of the imaged tumors, we show that this analysis is highly predictive of protein expression and that specific types of nanobubble time-intensity curve are associated with upregulation of different metabolic pathways. We applied this method to study the effects of two proteins, EphB4 and ephrinB2, which control tumor angiogenesis through bidirectional juxtacrine signaling, in mouse models of head and neck cancer. We show that ephrinB2 expression by endothelial cells and EphB4 expression by cancer cells have similar effects on tumor vasculature, despite sometimes opposite effects on tumor growth. This implicates a cancer-cell-intrinsic effect of EphB4 forward signaling and not angiogenesis in EphB4’s action as a tumor suppressor.

## Introduction

The *enhanced permeability and retention* effect is a phenomenon where nanoparticles accumulate in tumors as a result of increased permeability from vascular remodeling. This phenomenon has been widely documented^1,2^ and applied to delivery of chemotherapeutic agents to tumors using nanoparticles^3^. While vascular endothelial gaps are normally less than 7 nm, tumor blood vessels are thought to contain gaps in the range of 380-780 nm^4^, motivating the expectation that nanoparticles having diameters smaller than this should leak from tumor vasculature^5^. *Normalization* of tumor vasculature by blocking VEGFR-2 has been found to affect delivery of nanoparticles to tumors in a size-dependent manner, increasing delivery of smaller (12 nm) nanoparticles, while decreasing accumulation of larger (125 nm) nanoparticles, and this finding was attributed to decreased pore sizes^6^. However, this paradigm has recently been brought into question, with evidence suggesting that these endothelial cell gaps are very infrequent, and the majority of nanoparticles are actively transported *through* the endothelial cells, as opposed to going between them^7^.

Ultrasound contrast agents (UCAs) are typically 1-10 µm *micro*bubbles and are restricted to the microvasculature. The possibility of manufacturing stable, echogenic *nano*bubbles in the submicron range has been controversial^8^. Despite being substantially less echogenic, recent advances in shell composition have allowed for successful applications of nanobubble UCAs imaging in mice^5,9-11^. Nanobubble extravasation has been observed in mouse models of cancer^5,11-13^. The contrast signal from nanobubbles was found to decay more slowly than conventional microbubbles, and fluorophore-labeled nanobubbles were found to be more likely to extravasate, as measured by histology^11^. Nanobubble kinetics, described by contrast enhanced ultrasound (CEUS) time-intensity curves (TICs), have been found to differ from kinetics of conventional microbubbles^11^, and from the kinetics of nanoparticle contrast agents used in magnetic resonance imaging (MRI) and computed tomography (CT)^13^. Recent studies^12,13^ describe a “second-wave phenomenon” in nanobubble kinetics, which has not been observed with other UCAs or imaging modalities.

Eph receptors (named for the **e**rythropoietin-**p**roducing **h**epatocellular carcinoma cell line in which the first Eph gene was identified^14^) are the largest family of receptor tyrosine kinases (RTKs), consisting of 14 receptors in humans^15^, and bind to their membrane-bound *ephrin* ligands. Many Ephs have significant binding affinities for multiple ephrins and vice versa. Eph/ephrin signaling guides embryonic development, with expression of different receptors and ligands mediating cellular repulsion, migration, boundary formation, and axon guidance by inducing rearrangements of the cytoskeleton^16,17^. Some Eph/ephrins continue to be expressed in adult tissue, with ephrinB2 and EphB4 notably marking arterial or veinous identity blood vessels, with ephrinB2 being primarily expressed by arteries and EphB4 primarily by veins^18^. EphrinB2 has been shown to play a critical role in VEGF signaling, mediating VEGFR-2^19^ and VEGFR-3^20^ receptor internalization, necessary for activation of downstream signaling in VEGF-induced angiogenesis. By applying a novel CEUS image analysis method to mouse models with disrupted EphB4/ephrinB2 signaling, we were able to shed light on the complex roles of these proteins in tumor progression.

## Results

### Conventional CEUS image analysis reveals less persistent nanobubble signal in tumors lacking EphB4 or ephrinB2

We have previously shown that deletion of ephrinB2 (encoded by the *EfnB2* gene) from both cancer cells and endothelial cells has significant effects on the vasculature in mouse models of head and neck cancer, with decreased branching, improved perfusion, and increased pericyte coverage, associated with vascular normalization in mouse models of HNSCC^21^. EphB4 knockdown on cancer cells also increased VEGF-A expression^21^, and accelerated tumor growth. EphrinB2 is known to modulate angiogenesis by inducing internalization of VEGFR-2^19^ (**Fig. 1A**). To investigate changes in tumor vasculature detectable by CEUS, we performed a series of mouse experiments, using the MOC2 mouse oral squamous cell carcinoma cell line, where two experimental groups were compared to a common control (**Fig. 1B**). The first experimental group (ephrinB2-flox) uses EfnB2-flox/Tie2-Cre-ERT genetically engineered mice to inducibly delete ephrinB2 from endothelial cells, while the other two groups use wild-type C57BL6/J mice. The second experimental group (EphB4-ko) uses cancer cells where EphB4 has been knocked out using CRISPR (EphB4-ko), whereas the other two groups use control cells that were subjected to the same CRISPR selection process, but still express EphB4, as in Bhatia, et. al^21^.

**Fig. 1:**
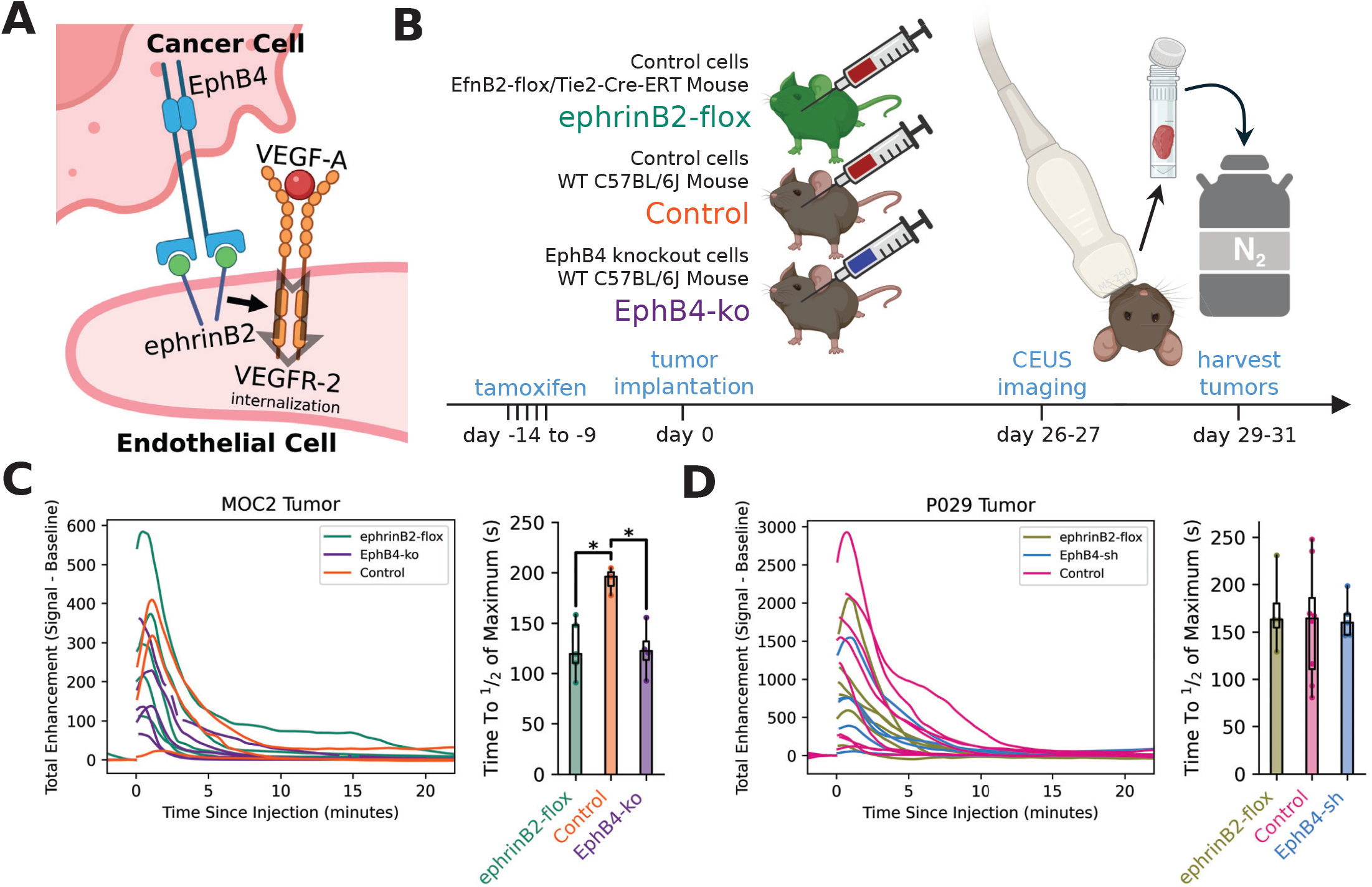
Effects of EphB4/ephrinB2 signaling on whole-tumor nanobubble kinetics. (**A**) Mechanism for effects of EphB4/ ephrinB2 signaling on tumor angiogenesis. (**B**) Timeline of MOC2 mouse experiments. (**C**) Time-intensity curves for MOC2 tumors using manual ROI selection in VEVO CQ (left) and time for contrast signal to wash out to ½ of its maximum value (right). Kruskal-Wallis test and Dunn pairwise comparison were statistically significant (p<0.05). (**D**) Time-intensity curves for P029 tumors using manual ROI selection in VEVO CQ (left) and time for contrast signal to wash out to ½ of its maximum value (right).

We initially performed imaging and analysis in a similar manner to Ramirez et. al^22^, obtaining one whole-tumor time-intensity curve for each mouse (**Fig. 1C**). Despite variability within groups, nanobubbles appeared to wash out of both ephrinB2-flox and EphB4-ko tumors faster than control tumors. Similar nanobubble washout in these groups contradicted our original hypothesis that vascular ephrinB2 and cancer cell EphB4 would have opposite effects on angiogenesis. These data instead suggested a simpler mechanistic explanation, that cancer cell knockdown/out of EphB4 primarily affects tumor angiogenesis by reducing the availability of binding partners for ephrinB2 reverse signaling—the mechanism depicted in **Fig. 1A**—and not by increasing VEGF production. To validate these results in a second model, we performed similar experiments using a genetically different cell line (P029), and knocked down EphB4 using sh-RNA instead of CRISPR, and treated with radiation therapy (XRT), which reduced tumor volumes at the time of imaging (**Supp. Fig. 1A**). We observed highly variable time-intensity curves, but no clear differences between groups (**Fig. 1D**).

A notable limitation of this analysis was that many time intensity-curves appeared to have shapes influenced by tissue with high intrinsic nonlinear signal moving into or out of the region of interest. To address this and reduce artifacts for further analysis, time-intensity curves were de-identified and blindly excluded or truncated (**Supp. Fig. 1B**). As a means of quantifying permeability of the tumor vasculature to nanobubbles, we propose a metric based on dividing areas under the time intensity curves (AUCs), where signal measured for the first 10 minutes is interpreted as proportional to the number of opportunities for a nanobubble to extravasate. Signal measured 15-20 minutes after injection is then interpreted as proportional to the number of nanobubbles that have extravasated into the interstitial fluid (**Supp. Fig. 1C**). In both models, we observed a trend for the fraction of nanobubbles extravasated to be lower in ephrinB2-flox than control tumors (**Supp. Fig. 1D**,**E**).

### Pixel clustering allows more detailed analysis of nanobubble kinetics

To more flexibly analyze the ultrasound data, we developed a Python library to work more efficiently with the raw pixel data generated from these experiments. Images were acquired roughly in 15-second cineloops at 30 frames per second started every 30 seconds, and some frames were gated out to reduce effects of breathing, resulting in many short periods of time missing from the data sets (**Fig. 2A**). To ease the process of excluding regions where large tissue-intrinsic nonlinear signals appear to vary over time (due to slight motion of the mouse during imaging), and directly address the fact that different regions of each tumor appeared to have very different nanobubble kinetics, we developed a pixel-clustering-based image segmentation method that classifies different regions of each tumor into “TIC clusters.” First, a large region of interest was manually selected for one tumor, then pixels within that region of interest were clustered into small groups of about 30 pixels with similar locations and nanobubble kinetics (**Fig. 2B**). A smoothed, interpolated, down-sampled, normalized time-intensity curve was then generated for each group of pixels. The time-intensity curves across all tumors were used for a second level of clustering (**Fig. 2C**). TICs were grouped into 12 clusters (**Supp. Fig. 2A**), but we limited our analysis to the largest 7 given that some of the less common clusters (e.g. small cluster 5) have median TICs with shapes that would be more plausibly represent changes in background intensity than nanobubble concentration. Somewhat similar analyses have been previously described^23,24^, but have not been used in a similar way, to the best of our knowledge.

**Fig. 2:**
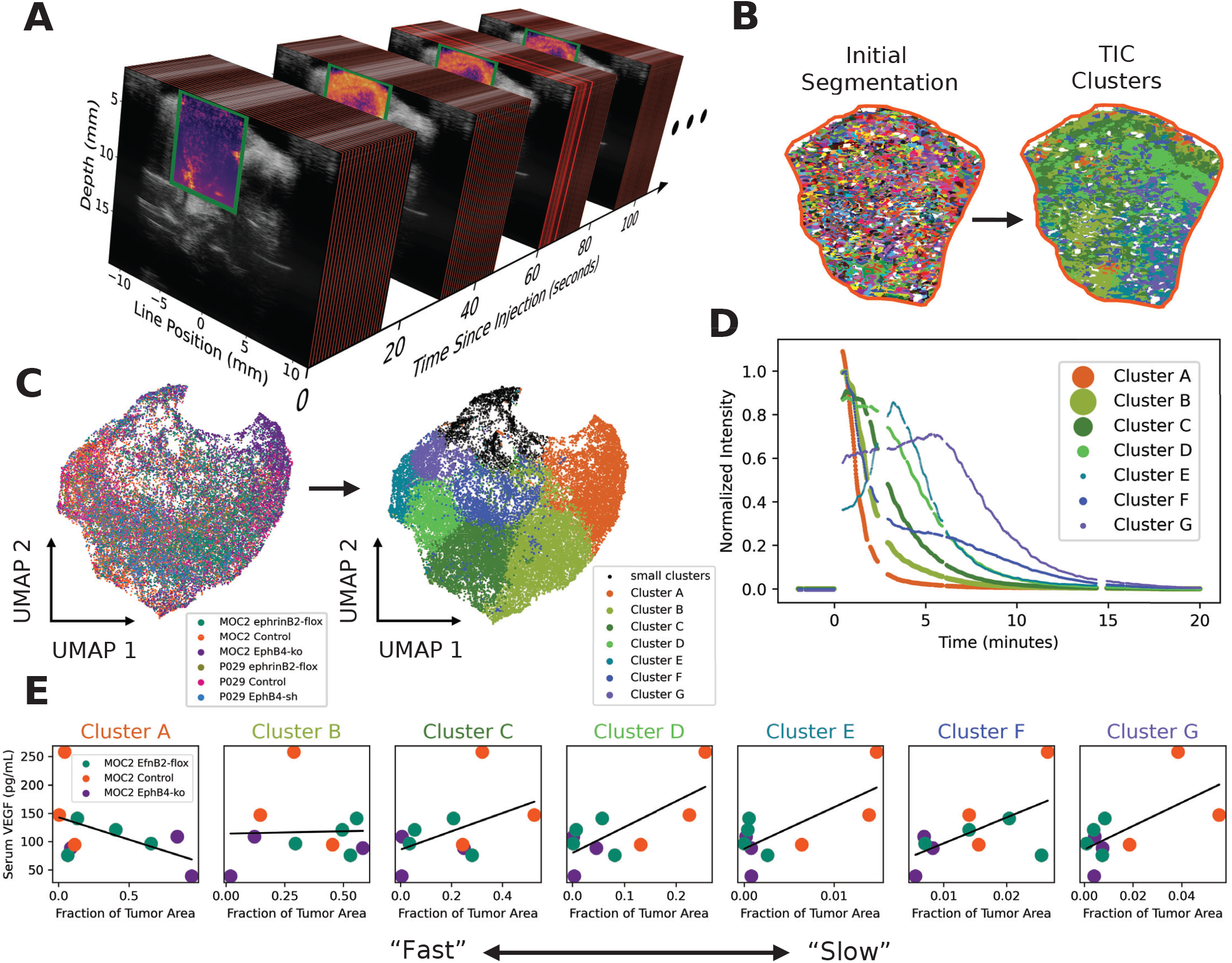
Nanobubble CEUS image segmentation by clustering of pixel-wise time intensity curves. (**A**) Graphical representation of missing data in the series of images acquired for one mouse. The larger grayscale image is B-mode used to locate anatomical features, the green rectangle indicates the smaller region for which nonlinear contrast data were acquired, and red represents frames that were excluded. (**B**) Representative image of initial pixel clustering and region classification segmentation. (**C**) UMAP plot of TICs for approximately 30-pixel regions of all tumors colored by group (left) and by K-Means clustering of the TICs (right). (**D**) Median TIC for each cluster colored in **C**. (**E**) Scatter plots of VEGF measured by ELISA of serum collected 8-10 days post implantation with correlation lines (black).

### Slow nanobubble wash-out is associated with elevated VEGF

Because we anticipated that differences in nanobubble kinetics would be partially controlled by VEGF-induced angiogenesis, we collected serum at multiple timepoints prior to ultrasound imaging and measured VEGF-A concentrations by ELISA. In serum samples collected 8-10 days post tumor implantation, VEGF correlated negatively with the fraction of tumor area classified as cluster A, the *fastest* TIC cluster, and positively with the slower clusters (**Fig. 2E, Supp. Fig. 2B**). As a measure of overall nanobubble wash-out delay, we computed the first moment of the mean TIC for each tumor, and also observed a trend for this to correlate positively with serum VEGF (**Supp. Fig. 2C**). Similar trends were also present for serum samples collected 20-days post implantation, approximately one week prior to imaging (**Supp. Fig. 2D**).

Correlations between serum VEGF and some of the excluded TIC clusters (**Supp. Fig. 2A**) were also observed, supporting the conclusion that more dramatically delayed nanobubble accumulation occurs in some regions. Considering each group individually, the very slow (excluded) TIC clusters correlated positively with serum VEGF only in the ephrinB2-flox group, while more common intermediate clusters correlated positively only in the control group, and no clear trend was observed in the EphB4-ko group (**Supp. Fig. 2E**). This is consistent with disruption or EphB4/ephrinB2 signaling leading to disruption of VEGF signaling (**Fig. 1A**).

### Cancer cell knockdown of EphB4 and deletion of endothelial ephrinB2 have similar effects on nanobubble kinetics, despite differences in protein expression

Tumor composition, in terms of nanobubble kinetics, differed between the MOC2 groups with three of the “slower” clusters, D, E, and G, being significantly less common in both ephrinB2-flox and EphB4-ko tumors than control (**Fig. 3A**). EphB4 knockdown tumors were also found to contain a lower density of blood vessels than control by immunofluorescence staining (**Fig. 2B,C, Supp. Fig. 3**). This supports the conclusion that cancer cell expression of EphB4 affects the vasculature primarily by stimulating ephrinB2 reverse signaling in endothelial cells, increasing angiogenesis.

**Fig. 3:**
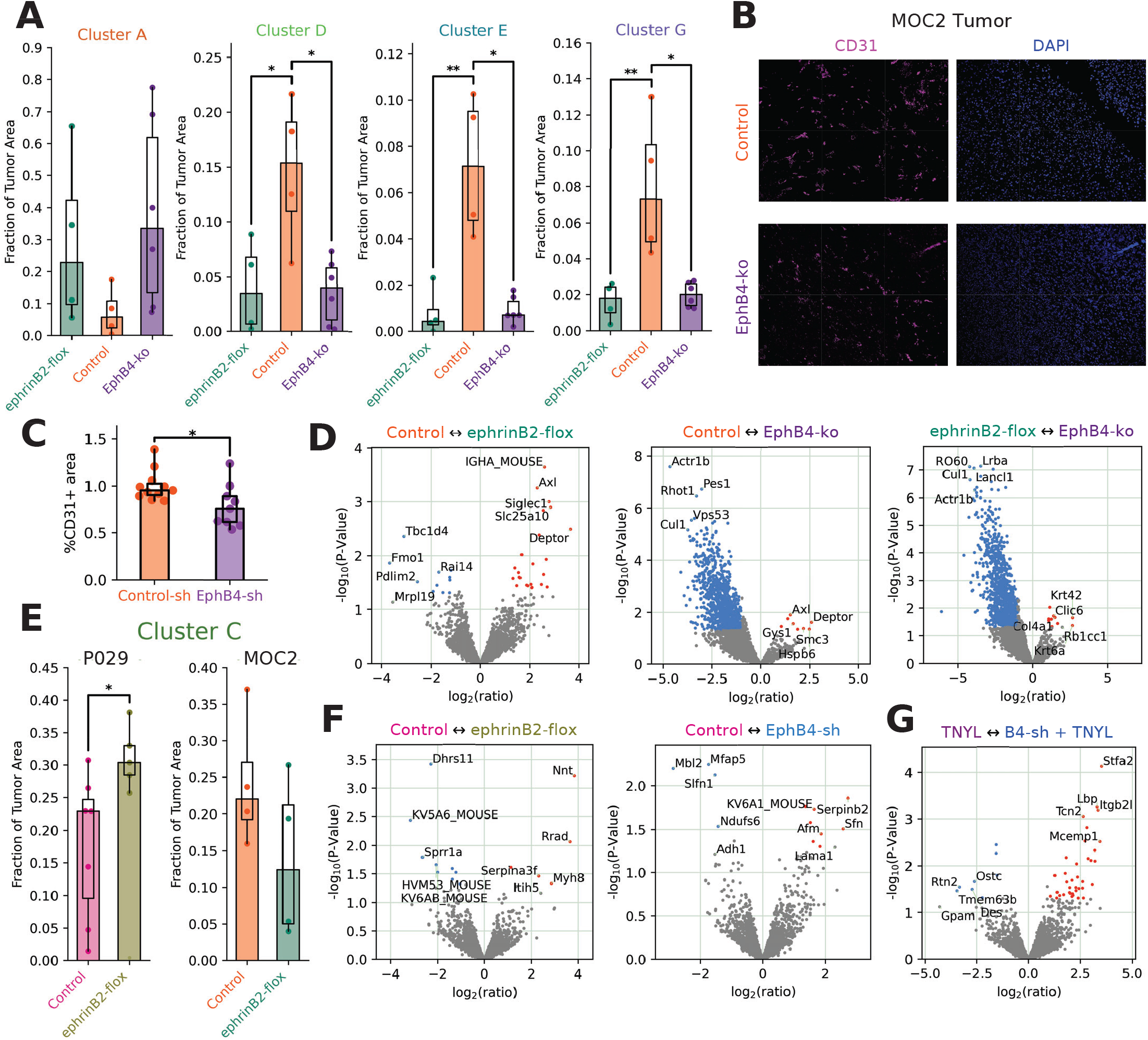
Deletion of ephrinB2 from endothelial cells or EphB4 from cancer cells have similar effects on vasculature. (**A**) Fractions of the area of each MOC2 tumor assigned to each of four selected TIC clusters. A Kruskal-Wallis test followed by Dunn pairwise comparisons was performed for each TIC cluster. In the comparison annotations, “*” indicates p < .05 and “**” indicates p < .01. (**B**) Representative immunofluorescence images of MOC2 Control and EphB4-sh tumors. (**C**) Vessel density (quantification of B, see **Supp. Fig. 3**). Student’s t-test comparing EphB4-sh to Control-sh gives p=.024. (**D**) Volcano plots of differential expression of cellular proteins between MOC2 groups. The “↔” in each plot title indicates that proteins on the left side of the plot are upregulated in the group to the left and vice versa. (**E**) Fractions of tumor area assigned to cluster C in control and EfnB2 flox mice. The “*” indicates p < .05 for a Mann-Whitney U test. (**F**) Volcano plots of differential expression of cellular proteins between CEUS-imaged P029 groups. (**G**) Volcano plot of differential expression of cellular proteins between TNYL-RAW-Fc treated P029 groups.

To further characterize differences between MOC2 groups, tumors were collected for proteomics approximately 3 days after CEUS imaging. Proteins from each sample were further separated into cellular, soluble extracellular matrix (ECM), and insoluble ECM fractions as in McCabe et. al^25^. Many cellular proteins were differentially expressed between MOC2 EphB4-ko and control, while few differences were observed between ephrinB2-flox and control (**Fig. 3D**). Some effects, including upregulation of Deptor, were shared in both comparisons and likely represent effects of endothelial ephrinB2 reverse signaling.

To determine if the observed effects were unique to MOC2, we applied the same analysis to the P029 data set. In P029, no statistically significant differences in TIC cluster composition were detected between groups using the Kruskal-Wallis test, but pairwise Mann-Whitney does show more cluster 1 area in ephrinB2-flox tumors than control, opposite to the trend observed in MOC2 (**Fig. 3E**). Single-sample GSEA of all tumors from the CEUS-imaged groups shows that different pathways tend to be up- and down-regulated in each tumor without obvious differences between groups (**Supp. Fig. 4**). While few individual cellular proteins were differentially expressed between P029 groups (**Fig. 3F**), GSEA using the fold-change-sorted protein list revealed several significantly upregulated pathways in EphB4-sh tumors, primarily related to immune response (**Supp. Fig, 5A**). Because the P029 tumors were treated with 8Gy × 3 fractions XRT, MOC2 tumors treated with the same radiation schedule were also included for comparison. GSEA showed many significant effects of EphB4-ko in MOC (**Supp. Fig. 5B**), but not on the same pathways as P029, underscoring that EphB4 plays a very different role in each model. Interestingly, both P029 and irradiated MOC2 groups had more differentially expressed proteins in the ECM components, while unirradiated tumors did not. These data point to more consistent effects on fibrosis in the context of XRT, which are not observed in the cellular component.

**Fig. 4:**
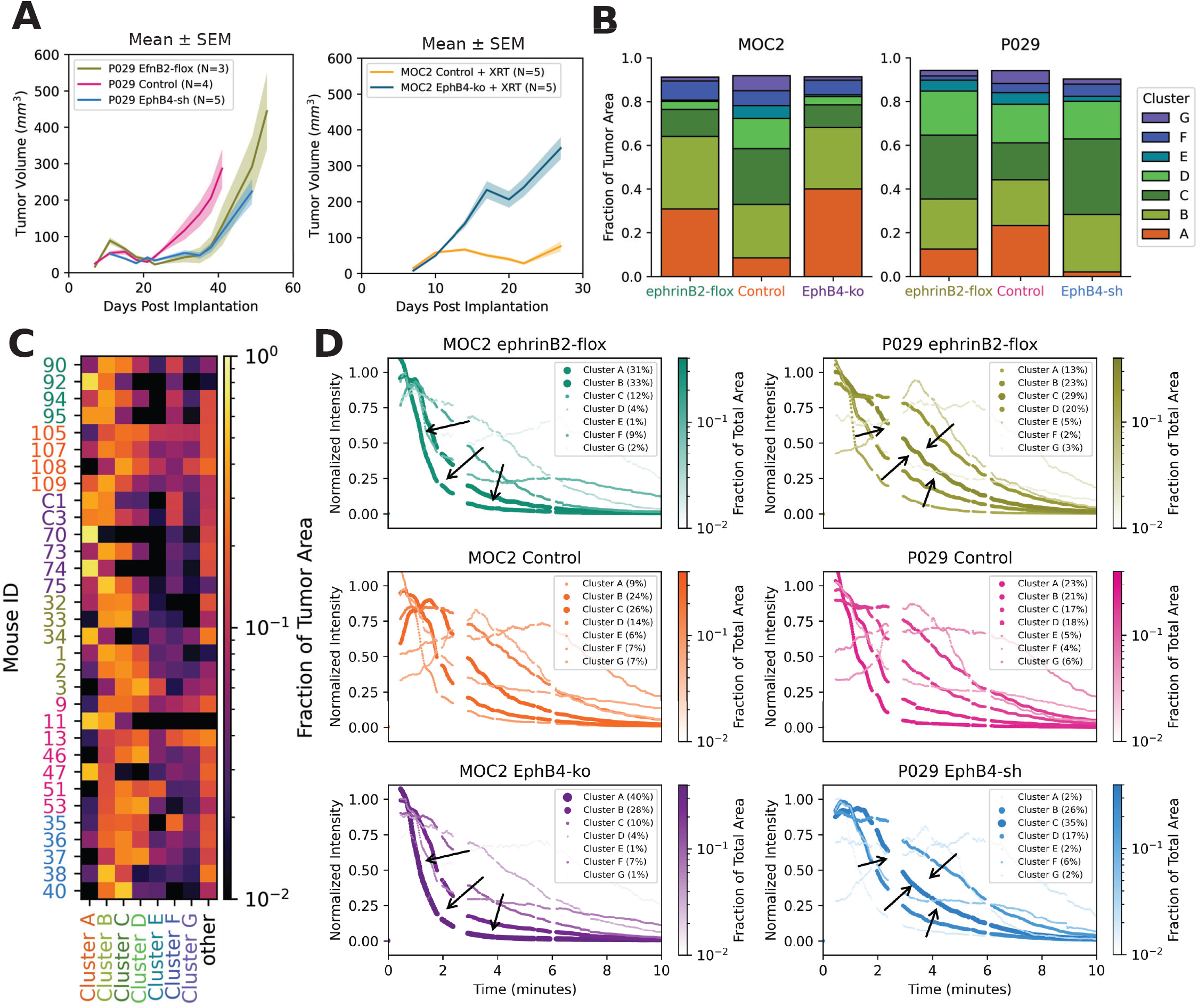
In a models where EphB4 KD opposite effects on tumor growth, it also has opposite effects on nanobubble kinetics. (**A**) Volumes of MOC2 and P029 tumors treated with 8Gy x 3 XRT, measured by automatic CBCT segmentation. Shaded regions indicate standard error on the mean. (**B**) Stacked bar chart of average area fractions for each TIC cluster in each group for MOC2 model (left) and P029 model (right). (**C**) Heatmap of TIC cluster compositions for each individual mouse. (**D**) Average TICs for each cluster within each group. The width of each line is proportional to the average image area fraction.

**Fig. 5:**
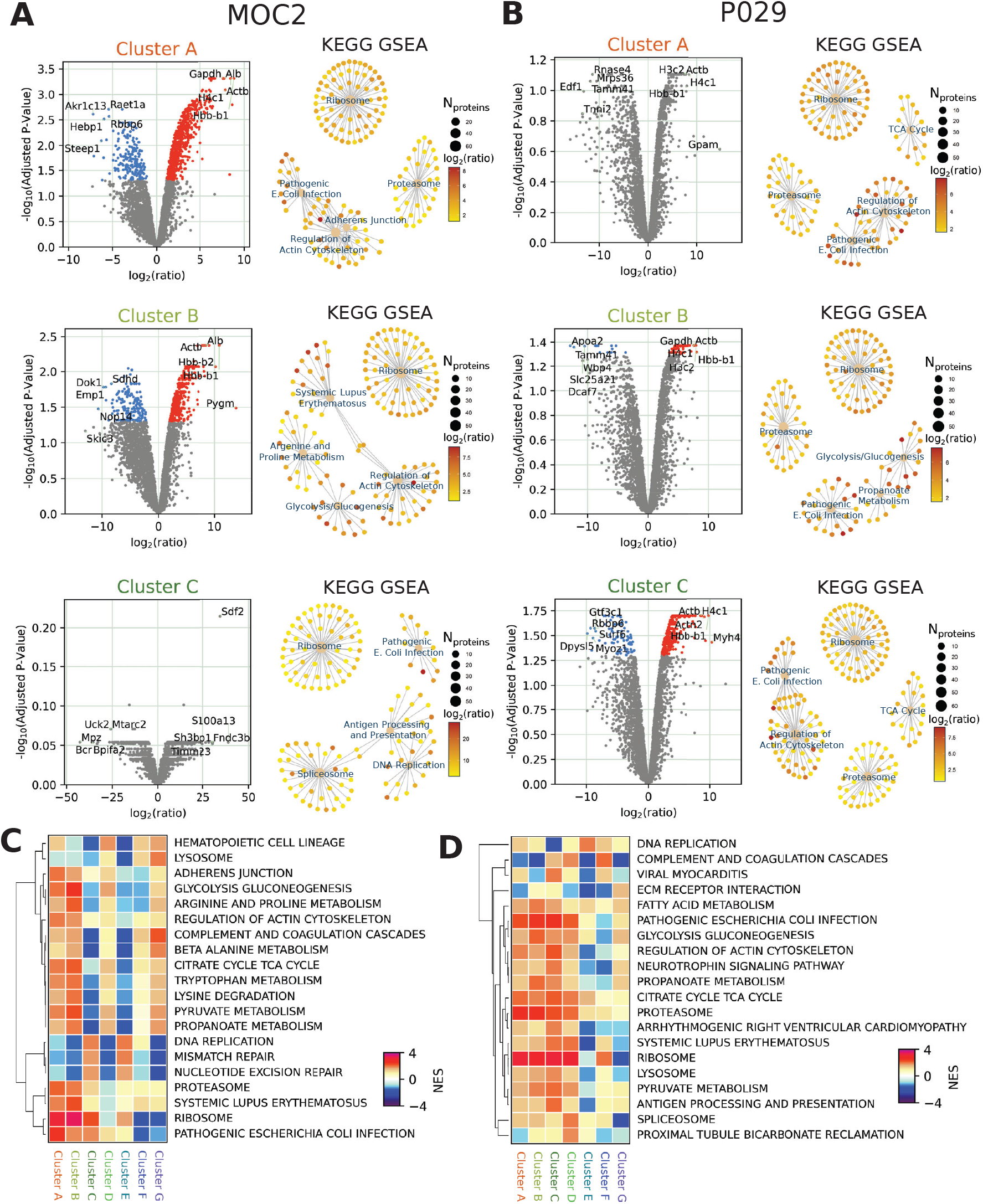
The most common TIC clusters are highly predictive of protein expression. (**A**) Volcano plots of protein expression associated with TIC clusters A, B, and C in Limma linear model controlling for group (left) and network plot of top 5 enriched KEGG pathways associated with each TIC cluster by GSEA (right) for MOC2. All pathways are significantly upregulated (adjusted p < .05). Labeled tan dots represent pathways and other dots represent proteins. (**B**) Volcano plots of protein expression associated with TIC clusters A, B, and C in Limma linear model (left) and network plot of top 5 enriched KEGG pathways associated with each TIC cluster by GSEA (right) for P029. All pathways are significantly upregulated (adjusted p < .05). Labeled tan dots represent pathways and other dots represent proteins. (**C**) Heatmap of normalized enrichment scores for the top 20 KEGG pathways ranked by maximum absolute value for any TIC cluster in MOC2 tumors. (**D**) Heatmap of normalized enrichment scores for the top 20 KEGG pathways ranked by maximum absolute value for any TIC cluster in P029 tumors.

To dissect vascular effects from the cancer cell effects in the P029 model, we employed a pharmacological inhibitor in two additional groups. Treatment with TNYL-RAW-Fc plasmid (**Fig. 2G**), which induces sustained production of a peptide that inhibits EphB4-ephrinB2 interaction^26^, has previously been shown to reduce tumor growth in models where EphB4 cancer cell knockdown accelerates it^27^, suggesting that the drug acts on EphB4 expressed by other cells. Consistent with this hypothesis, TNYL-RAW-Fc had more pronounced effects on protein expression in P029 EphB4-sh tumors (**Fig. 2G, Supp. Fig. 5B**). Clearly, EphB4’s effect on angiogenesis is only one aspect of a more complex role in shaping the tumor microenvironment.

### EphB4 knockdown has opposite effects on both tumor growth and nanobubble kinetics in P029 compared to MOC2 tumors

Contrary to previous findings^21^, knockdown of EphB4 in the P029 model reduced tumor growth rather than accelerating it. To verify that this was driven by cancer-cell-intrinsic effects of EphB4 in P029 and was not an effect of radiation therapy, we treated both MOC2 and P029 tumors with the same 8Gy × 3 fractions XRT at 8, 11, and 14 days post implantation. We monitored tumor growth by automated segmentation of CBCT scans^28^ to provide robust tumor volume measurements. While EphB4 cancer cell knockout markedly reduced response to radiation in MOC2, both EphB4-sh and ephrinB2-flox similarly delayed tumor regrowth in P209 (**Fig. 4A**). While both cell lines harbor Kras mutations^29,30^, these data suggest that proliferation or cell death may be driven by very different pathways downstream of EphB4.

In both MOC2 and P029 models, removing EphB4 from cancer cells and removing ephrinB2 from endothelial cells had similar effects on the prevalence of each TIC cluster (**Fig. 4B**). However, the specific changes were different, with Cluster A being more common in MOC2 ephrinB2-flox and EphB4-ko, while the opposite was true for P029. Similarly, in the P029 experimental groups, cluster C was more common, while the opposite was true for MOC2. All groups display substantial variability between individual tumors (**Fig. 4C**), but the overall trends paint a clear picture of how disruption of EphB4/ephrinB2 signaling tends to affect nanobubble kinetics in each model. To visualize this in terms of TIC shapes, we plotted the median TIC for each cluster within each group with line thickness proportional to the average fraction of tumor area classified into each cluster (**Fig. 4D**). In general, it appears that inhibition of endothelial ephrinB2 signaling yields faster nanobubble wash-out in MOC2, but slower wash-out in P029.

### The most common TIC clusters are highly predictive of similar protein expression in both MOC2 and P029

To further explore the biological significance of differences in nanobubble kinetics, we used the R package Limma to fit linear models and identify protein signatures associated with each cluster, while controlling for groups as covariates. Statistically significant associations with individual proteins were observed for the most common TIC clusters (**Fig. 5A,B**, left columns). Many of the same proteins, including hemoglobin, were positively associated with all 3 clusters. GSEA analysis of proteins ranked in this way using KEGG pathway gene sets (**Fig. 5A,B**, right columns) shows that ribosome, proteasome, and several metabolic and immune response pathways are positively associated with these types of nanobubble kinetics in both MOC2 and P029. These might be interpreted as characteristic of tumors with more “normal” vasculature in contrast to the “slow” clusters (E, F, and G) representing a less common phenotype with reduced blood flow.

We computed GSEA results for all clusters and made heatmaps of the normalized enrichment scores to compare pathways associated with each TIC cluster (**Fig. 5C**,**D**). This revealed distinct phenotypic groupings of the TIC clusters. Specifically, clusters A and B in both models and clusters C and D in P029 are associated with a similar metabolically active phenotype, with upregulated glycolysis and TCA cycle. However, cluster C was associated with downregulation of these pathways in MOC2, suggesting that a vascular phenotype with somewhat slower nanobubble wash-out was conducive to ATP production in P029, but not MOC2 tumors. This is particularly interesting in light of the opposite effects of ephrinB2 signaling on prevalence of this cluster in the two models (**Fig. 3E**).

Results for the slower TIC clusters (E, F, and G) were less consistent. These time-intensity curves were more aberrant and may each be associated with distinct types of tumor vasculature. Clusters E and G represent regions where the nanobubble signal peaks several minutes after injection, likely because they are not in close proximality to large, functional blood vessels, and cluster F is approximately a linear combination of clusters A and G (**Fig. 2D**), which might be due to the signal from a larger vessel detected in a nearby hypovascular region. Because these clusters represent only small fractions of the imaged plane, and therefore small fractions of the protein analyzed, we reasoned that proteins detected as up- or down-regulated in tumors containing more of these TIC types may be produced elsewhere. Statistically significant up- and down-regulation of various pathways was observed for all seven TIC clusters for both MOC2 (**Supp. Fig. 7**) and P029 (**Supp. Fig. 8**) tumors using GO Biological process gene sets. This is particularly impressive in light of the highly variable single-sample GSEA results within each group (**Supp. Fig. 4**). Collectively, these results demonstrate that nanobubble kinetics can be used to predict biological functions in tumors far beyond the effects of manipulating EphB4 and ephrinB2 expression.

### Heterogeneity of nanobubble kinetics is reduced by removal of ephrinB2 from endothelial cells or EphB4 from cancer cells

To ground the interpretation of these correlations, we examined how the tumors were spatially segmented to derive fractions of area classified into each cluster (**Fig. 6A,B**). Because TICs fill a high dimensional space without clear boundaries between clusters (**Fig. 2C**), small scale mixing of clusters A and B may occur in regions with fairly uniform nanobubble kinetics that are intermediate between the centers of those clusters. MOC2 ephrinB2-flox and EphB4-ko tumors tended to contain large areas with no more than two TICs clusters, in contrast to control tumors, which are more heterogeneous. While some tumors heavily favored cluster A over cluster B or vice versa, uniformity was a more consistent feature of these groups.

**Fig. 6:**
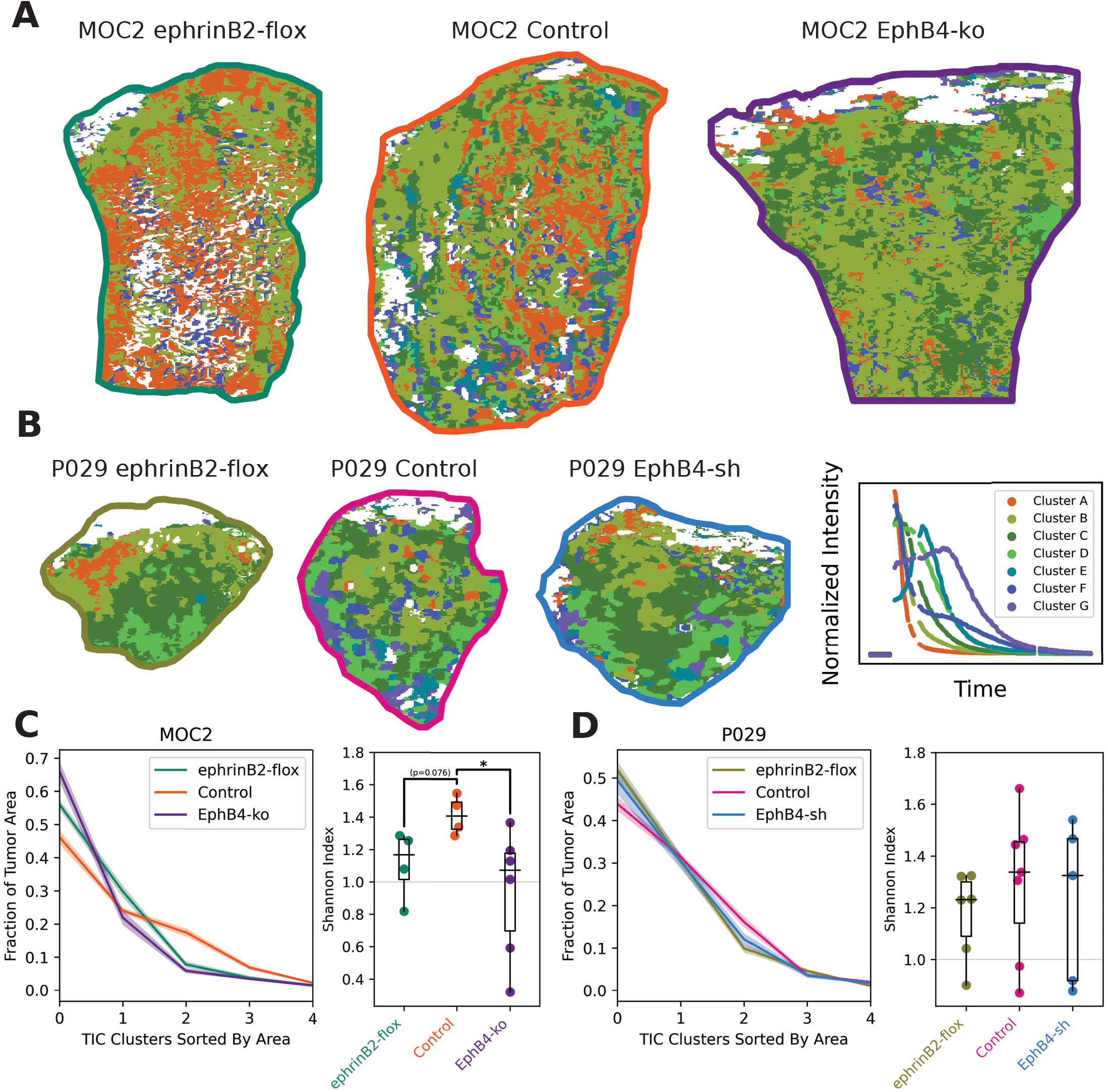
Removal of ephrinB2 from endothelial cells or EphB4 from cancer cells reduces intertumoral heterogeneity of nanobubble kinetics. (**A**) Representative pixel clustering segmentations of one MOC2 tumor from each group. White represents regions that either fell into one of the excluded TIC clusters (see **Supp. Fig. 2A**) or were excluded from clustering because of saturated pixels. (**B**) Representative pixel clustering segmentations of one P029 tumor from each group. These and the MOC2 images above are all identically scaled. A smaller reproduction of **Fig. 2D** is included to the right for easier interpretation of the region colors. (**C**) Sorted label area trends with shaded regions indicating standard error on the mean (left) and Shannon index quantification (right) for MOC2 tumors. A Kruskal Wallis test gave (p=.0506), and Dunn pairwise comparisons were performed (taking α=.1) to generate the p-values indicated above the comparison bars. (**D**) Sorted label area trends with shaded regions indicating standard error on the mean (left) and Shannon index quantification (right) for P029 tumors.

To further quantify the degree of heterogeneity, we sorted the TIC cluster areas within each tumor and plotted the mean and standard error on the mean for each group (**Fig. 6C**, left). We also computed the Shannon index of the cluster composition for each tumor (**Fig. 6C**, right), and found that this was higher in MOC2 control tumors than in either experimental group. Control tumors tended to have less of the most common TIC cluster and more of the third most common, indicating increased intratumoral heterogeneity. The same analysis applied to P029 tumors showed similar trends, but more variability within groups, perhaps because the tumors were smaller at the time of imaging (**Fig. 6D**).

## Discussion

A significant strength of the present study is the large number of mice imaged, capturing a range of tumors from multiple groups. To the best of our knowledge, no attempt has previously been made to similarly use nanobubble CEUS as a tool for cancer biology research. One obstacle to this research was the long imaging sessions required to fully observe nanobubble wash-out curves. This not only causes technical challenges for the image analysis but reduces the number of mice that can feasibly be imaged. This may adversely affect the applicability of this kind of imaging in human patients. It might, therefore, be advantageous to determine how much useful information is lost by imaging for significantly shorter periods of time.

Surprisingly strong correlations between TIC cluster composition and protein expression in these tumors may suggest that the single plane of each tumor imaged is representative of the rest of the tumor. However, 3D images^31^ including the whole tumor would likely give more accurate results and allow for 3D motion tracking, so that small features entering or exiting the imaged plane could not cause unexplained shifts in signal intensity. Particularly in P029 tumors, nanobubble kinetics tended to change with depth, with slower TIC clusters appearing near the tumor core. The core may be over-represented in extrapolating from fractions of the imaged plane to fractions of the full tumor volume in this analysis.

Correlations with bulk proteomics provide insight into biological processes associated with regional TICs, but spatial-omics technology could provide additional information. Further optimization of the CEUS image analysis protocol may provide a more robust analysis. Clustering TICs into asmall number of categories may be sub-optimal because doing so requires drawing arbitrary divisions between clusters that could more accurately be described by fitting parameters of a kinetic model. It may also be possible to integrate our clustering approach with other image analysis methods developed for nanobubble CEUS^32^.

The data presented here eliminate pro-angiogenic effects as a likely mechanism for the accelerated tumor growth associated with EphB4 cancer cell knockdown. We had previously shown that EphB4 knockdown cells proliferate faster in vitro^21^, which could result from decreased competition for ephrinB2 binding leading to increased forward signaling through another receptor or an unknown mechanism of tumor suppression downstream of EphB4. Additional experiments will be required to address this.

The potential impact of nanobubble technology extends far beyond studying the role of EphB4/ephrinB2 signaling in tumor angiogenesis. Nanobubbles are of interest as vehicles for drug delivery^32,33^, and understanding their unique, spatially heterogeneous kinetics could crucially inform predications of how effects of drugs delivered in this way will differ compared to other delivery methods. Targeted UCAs conjugated to antibodies^5,34^ have also been studied in this context. Image analysis that effectively accounts for intratumoral heterogeneity will be crucial for developing nanobubbles a tool for clinical and research applications.

## Materials and Methods

### Mouse Models

All animal procedures were conducted in accordance with protocols approved by the University of Colorado, Anschutz Medical Campus institutional animal care and use committee (IACUC). Some mice were euthanized due to tumor volumes exceeding 1000 mm^3^, weight loss exceeding 5% of body weight per day over 2-3 days, or abnormal breathing due to lung metastases. In rare cases, mice were euthanized without reaching these endpoints because of tumor ulceration or died unexpectedly after ultrasound imaging or blood collection.

Female C57BL/6 wild-type or EfnB2^fl/fl^Tie2-Cre-ERT (ephrinB2-flox) mice where implanted with MOC2^30^ or P029^29^ cells in the right buccal, as described in Oweida, et al.^35^. Five 1mg doses of tamoxifen were administered to ephrinB2-flox mice by IP injection >7 days prior to tumor implantation to induce vascular ephrinB2 knockout. Tamoxifen was also administered to wild-type mice to control for any potential effect of tamoxifen on tumor growth.

### Cell Lines

The MOC2 cell line was obtained from Dr. Ravindra Uppaluri (Dana-Farber Cancer Institute, Boston, MA)^21^, the P029 cell line from the Xiao-Jing Wang lab (University of Colorado, Anschutz Medical Campus). Genetic manipulations (to knock down/out EphB4) of the MOC2 and P029 cell lines were performed by the University of Colorado Cancer Center Functional Genomics Facility. MOC2 cells were transfected with PX458 control plasmid or PX458 containing gRNA targeting EPHB4 and CRISPR knockout (The same cells were used in Bhatia, et al., 2022)^21^. Knockdown/out of EphB4 was confirmed by western blot.

### Immunofluorescence

Paraffin-embedded MOC2 tumor slides were stained with anti-CD31 primary antibody (Cell Signaling 77699S) at 1:400 dilution and Alexa Fluor 647 secondary antibody (**A-31573**) at 1:400 dilution. Images were acquired using a Keyence BZ-X800 microscope at 20x magnification and analyzed in Python.

### Radiation Therapy

X-ray irradiation was delivered in three fractions of 8Gy at 8, 11, and 14 days after tumor implantation using the X-Rad SmART small animal irradiator (Precision X-ray Inc., Madison CT). Mice were positioned under isoflurane anesthesia and beams aligned using fluoroscopy. The treatment plan was simulated using SmART-ATP software (SmART Scientific Solutions, Maastricht, the Netherlands) with a CBCT image of a representative model mouse, then each tumor was irradiated for the same amount of time, based on a calculated dose rate of 4.7Gy per minute.

### Ultrasound Imaging

Ultrasound imaging was performed according to previously described methods^22^. Briefly, B-mode and nonlinear contrast images were acquired using the VEVO 2100 ultrasound machine with a frequency of 18 MHz at approximately 30 frames per second. Mice were positioned with isoflurane anesthesia under a heat lamp with a rectal thermometer to monitor body temperature and electrodes connected to fore- and hind-limbs to monitor breathing and heart rate during imaging (Fig. 1A). Due to a limit on the amount of nonlinear contrast data the VEVO 2100 can acquire at a time, data were collected in cine-loops of approximately 20 seconds at approximately 30 second intervals for 20 to 40 minutes, including 3-7 minutes of baseline data prior to injection of 120 µL of nanobubble contrast agent via tail vein catheter. The contrast agent was prepared for no more than two mice at a time to reduce differences in time from activation and isolation of the nanobubbles to imaging.

### Tumor Measurement and CBCT Imaging

CBCT scans were collected approximately twice per week using the X-Rad SmART small animal irradiator under isoflurane anesthesia. The tumor volumes reported were derived from automated segmentation of the CBCT scans, as previously described^28^.

### Nanobubble Preparation

Nanobubbles were prepared as in Perera, et. al.^36^ and Hernandez, et al.^37^, using lipid solutions obtained from the Exner lab at Case Western Reserve University (Cleveland, OH). Briefly, a lipid solution consisting of 6.1 mg DBPC (1,2-dibehenoyl-sn-glycero-3-phosphocholine), 2 mg DPPE (1,2-dipalmitoyl-sn-glycero-3-phosphoethanolamine), 1mg DPPA (1,2-dipalmitoyl-sn-glycero-3-phosphate) and 1mg mPEG-DSPE (1,2-distearoyl-sn-glycero-3-phosphoethanolamine-N-[methoxy (polyethylene glycol)-2000]) was prepared in 0.1 mL propylene glycol by repeated sonication and heating to 80°C. A second mixture of 0.8 mL PBS (phosphate buffered saline) and 0.1 mL glycerol, also heated to 80°C was then added to the lipid solution and sonicated at room temperature for 10 minutes. This solution was then transferred to a 3 mL vial with a rubber seal and the gas in the vial replaced with octafluoropropane (C_3_F_8_). The bubbles were activated by agitation in a mechanical shaker for 45 s, approximately 2 hours prior to ultrasound imaging, then centrifuged at 50*g* for 5 minutes to isolate submicron bubbles based on buoyancy. Nanobubbles were diluted 1:5 in PBS and approximately 200 µl of this solution was then pipetted into the back of an insulin syringe and used as the UCA (ultrasound contrast agent). A different vial was used for each mouse to mitigate potential effects of differences in nanobubble preparation on results.

### Ultrasound Image Analysis

Conventional quantification of region-of-interest (ROI)-averaged contrast signals were performed using VEVO LAB/VEVO CQ software (FujiFilm VisualSonics, Toronto, Canada). Additional processing and analyses, including background subtraction, gating, and smoothing were then applied to the resulting time-intensity curves in Python.

Pixel clustering analysis was implemented in Python using primarily NumPy, SciPy, and Scikit-learn. A region of interest corresponding to the tumor was first manually selected for each mouse using a custom graphical user interface defined in Python. Next, a vector including both a time-intensity curve and weighted x/y coordinates for each pixel position was used for k-means clustering within each tumor to obtain an initial segmentation as shown in the left panel of Fig. 1D. The number of regions per tumor was chosen to give an average area of 30 pixels for each region.

Time-intensity curves for each region were then down-sampled by taking the median pixel intensities for one-second time intervals starting two minutes before injection of the contrast agent and ending 20 minutes after. Regions with more than 5 saturated medians (having values of either 0 or 2^16^-1) were then excluded and smoothing interpolation was applied to the remaining down-sampled time-intensity curves to reduce noise and fill gaps between cine-loops. These curves were then linearly rescaled to have 1% of values less than zero and 1% of values greater than 1. A second level of k-means clustering was then applied to group the processed time-intensity curves for all mice into 12 clusters. The 5 smallest clusters, each representing less than 3% of regions, were excluded from most analyses **(Supp. Fig. 2A)**.

### Proteomics

Tumors were harvested, immediately flash-frozen in liquid nitrogen, and lyophilized prior to analysis. Preparation of tumor samples for mass spectrometry was performed as previously described^25,38^. Lyophilized samples were delipidated by successive extractions with 2 ml 100% ice-cold acetone and processed by a stepwise extraction with CHAPS and high salt, guanidine hydrochloride, and chemical digestion with hydroxylamine hydrochloride (HA) in Gnd-HCl generating cellular, soluble ECM (sECM), and insoluble ECM (iECM) fractions for each sample, respectively. Protein concentration of each fraction for each sample was measured using A660 Protein Assay (Pierce™). Thirty μg of protein resulting from each fraction was subjected to proteolytic digestion using a filter-aided sample preparation (FASP) protocol^39^ with 10 kDa molecular weight cutoff filters (Sartorius Vivacon 500 #VN01H02). Samples were reduced with 5mM tris(2-carboxyethyl)phosphine), alkylated with 50 mM 2-chloroacetamide, and digested overnight with trypsin (enzyme:substrate ratio 1:100) at 37ºC. Digested peptides were recovered from the filter using successive washes with 0.2% formic acid (FA), loaded onto Evotips, and analyzed directly using an Evosep One liquid chromatography system (Evosep Biosystems, Denmark) coupled to a timsTOF Pro mass spectrometer (Bruker Daltonics, Bremen, Germany) as previously described^40^.

Fragmentation spectra were interpreted against the UniProt mouse proteome database using the MSFragger-based FragPipe computational platform^41^. Contaminants and reverse decoys were added to the database automatically. The precursor-ion mass tolerance and fragment-ion mass tolerance were set to 15 ppm and 0.08 Da, respectively. Fixed modifications were set as carbamidomethyl (C), and variable modifications were set as oxidation (M), oxidation (P) (hydroxyproline), Gln->pyro-Glu (N-term), deamidated (NQ), and acetyl (Protein N-term). Two missed tryptic cleavages were allowed, and the protein-level false discovery rate (FDR) was ≤ 1%.

The resulting data were then analyzed in R. First, filtering and imputation were performed using the package DEP to handle missing values. Next, the package limma was used to detect proteins differentially expressed between groups or associated with specific TIC clusters by fitting linear models. Results from limma were then used to rank proteins for GSEA analysis using clusterProfiler. Single-sample GSEA analysis was performed using the package GSVA.

### Statistical Analysis

Statistical analyses were performed in Python. The stats SciPy module was used to calculate correlation coefficients and associated p-values and perform Kruskal-Wallis and Mann-Whitney tests. Dunn tests were performed using the package scikit-posthocs without correction for multiple comparisons. Benjamini-Hochberg-adjusted p-values for differentially expressed proteins were calculated in R by Limma.

## Acknowledgements

This study was supported in part by the National Institutes of Health P30CA06934 funded Mass Spectrometry Proteomics Shared Resource [RRID SCR_021988]. **Fig. 1A,C** use images from BioRender.com. The Karam laboratory would like to gratefully acknowledge support from NIH (R01DE028529, R01CA28465, R01DE028529, 1P50CA261605-01), and V Foundation (to SDK). The Benninger laboratory gratefully acknowledges support from NIH/NIDDK (R01DK106412, R01DK102950, R01DK140904, to RKPB), Juvenile Diabetes Research Foundation Grants 1-INO-2019-783-S-B and 3-SRA-2024-1554-S-B (to RKPB and AAE), and the University of Colorado Diabetes Research center (P30 DK116073), as well as NCATS (TL1 TR002533), and NIDKK (T32 DK120520, F31 DK136186) (to MC).

