## Supplemental Figures for "Heterogeneous Kinetics of Nanobubble Ultrasound Contrast Agent and Angiogenic Signaling in Head and Neck Cancer"

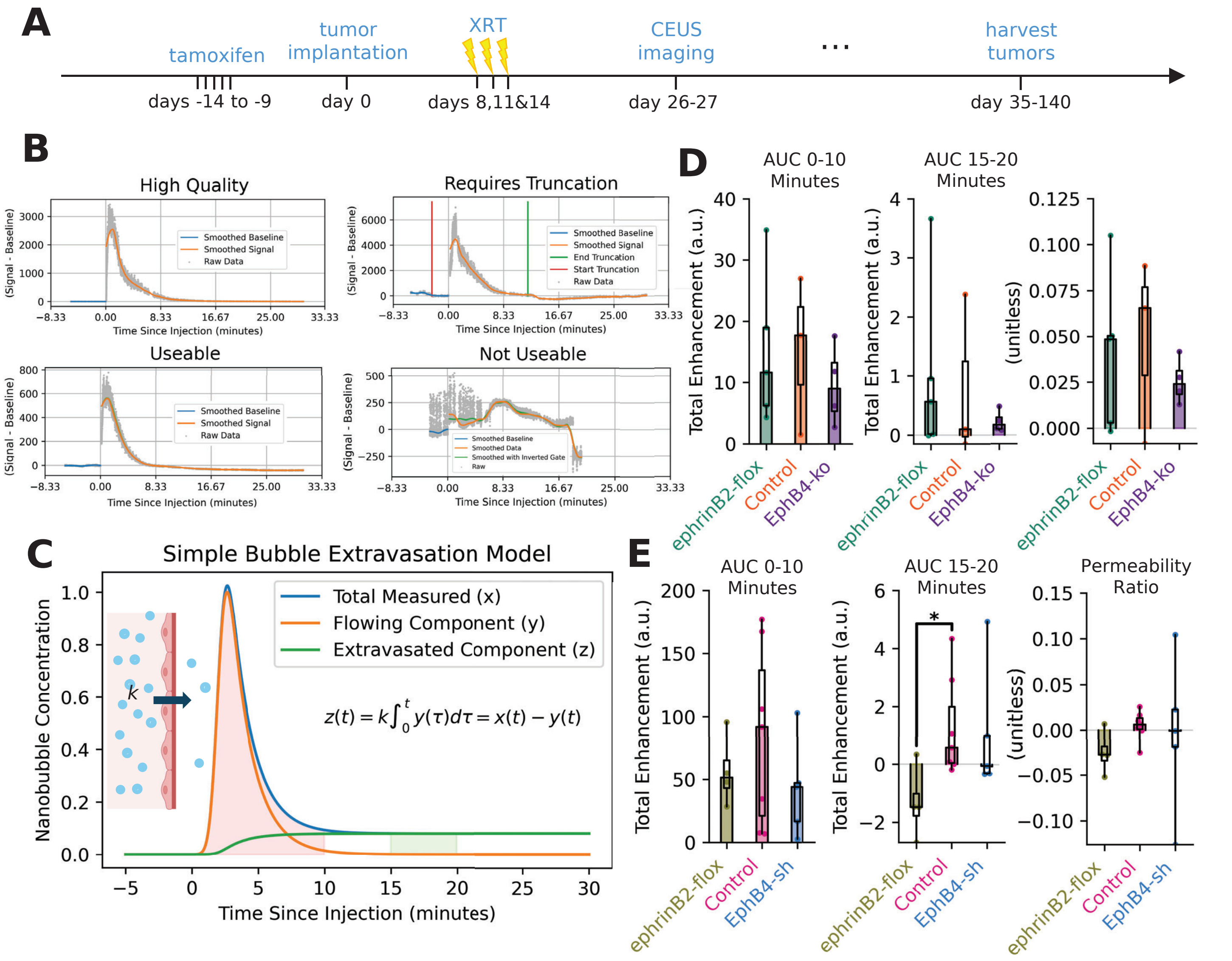


**Supp. Fig. 1: Vascular permeability estimation using manual ROI selection.**

(**A**) Timeline of P029 nanobubble experiments. CT scans were also acquired approximately every 4 days to monitor tumor growth. (**B**) Representative time-intensity curves for 4 categories of TIC used identified for blind manual data curation. (**C**) Theoretical plot (not real data) to explain how permeability of tumor vasculature to nanobubbles can be estimated from areas under the timeintensity curves. (**D**) AUCs used to estimate permeability of the vasculature to nanobubbles for MOC2 tumors (left, center) and ratio between them (right). (**E**) AUCs used to estimate permeability of the vasculature to nanobubbles for P029 tumors (left, center) and ratio between them (right). Kruskal-Wallis test and Dunn post-hoc test were performed. The p-value for the pair marked "*" is p=0.0216, with no adjustment for multiple comparisons.

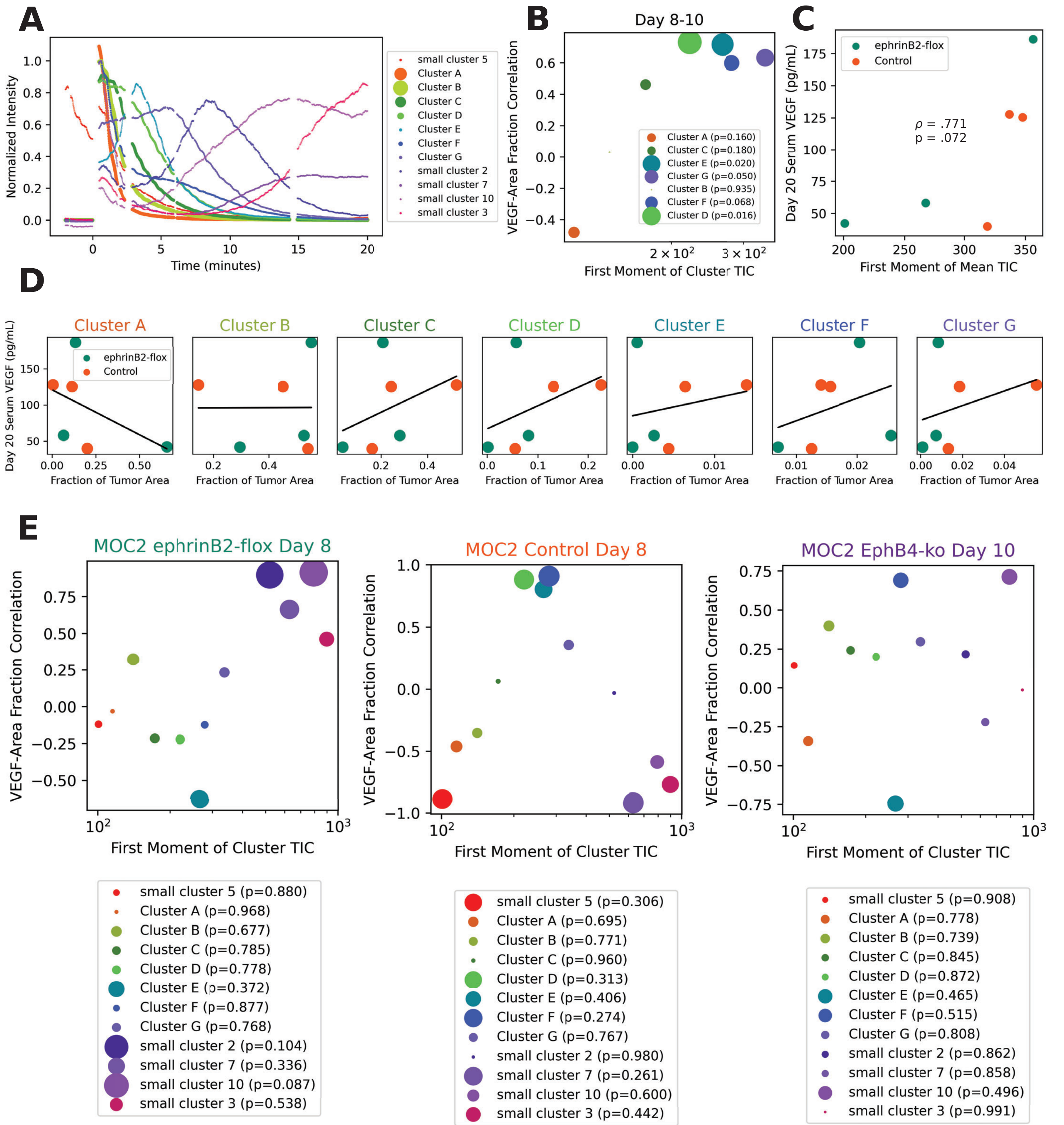


**Supp. Fig. 2: Serum VEGF from various timepoints correlate with nanobubble TIC cluster compositions.** (**A**) Median TICs for each cluster, including small clusters excluded from **Fig. 2D**. (**B**) Day 20 Serum VEGF plotted against first moment of whole-tumor mean TIC. Spearman correlation statistics are shown in the center. (**C**) VEGF measured by ELISA in serum collected 20 days post implantation plotted against faction of tumor area classified as each cluster with correlation lines (black). (**D**) Pearson's correlation coefficients between VEGF in serum collected 8-10 days post tumor implantation and tumor area assigned to each TIC cluster plotted against first moment of each cluster's mean TIC. (**E**) Scatter plots of Pearson's correlation coefficients between serum VEGF concentration and fraction of tumor area in each TIC cluster for the individual groups shown in **Fig. 2D**. Larger dots correspond to lower p-values.

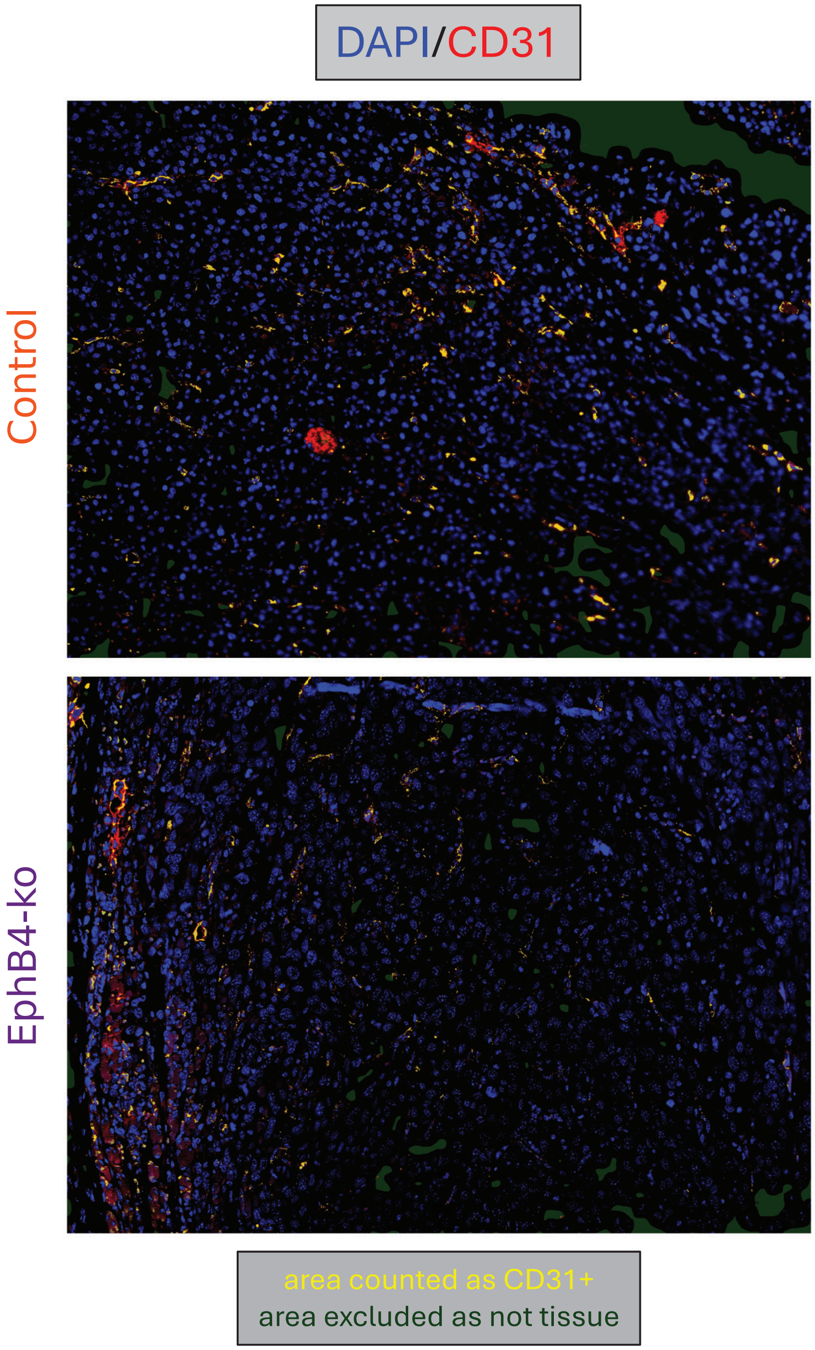


**Supp. Fig. 3: CD31 quantification for IF images.**
Representative immunofluorescence images used for vessel density quantification (**Fig. 3C**) showing DAPI and CD31 staining with additional colors to explain area percentage quantification. Green represents areas excluded as not containing tissue, due to lack of DAPI staining. Yellow represents area classified as CD31+. Both image channels were linearly rescaled to have 70% black and 1% maximum-brightness pixels. A blurred version if the resulting DAPI image was then thresholded to detect green excluded regions. To detect, blood vessels, the DAPI channel was then subtracted from the CD31 channel to reduce effects of folded tissue. Two copies of the CD31 channel blurred by different amounts were then compared to exclude areas with relatively flat high values in the red channel, likely due to non-specific staining. The same steps were applied identically to all images in Python.

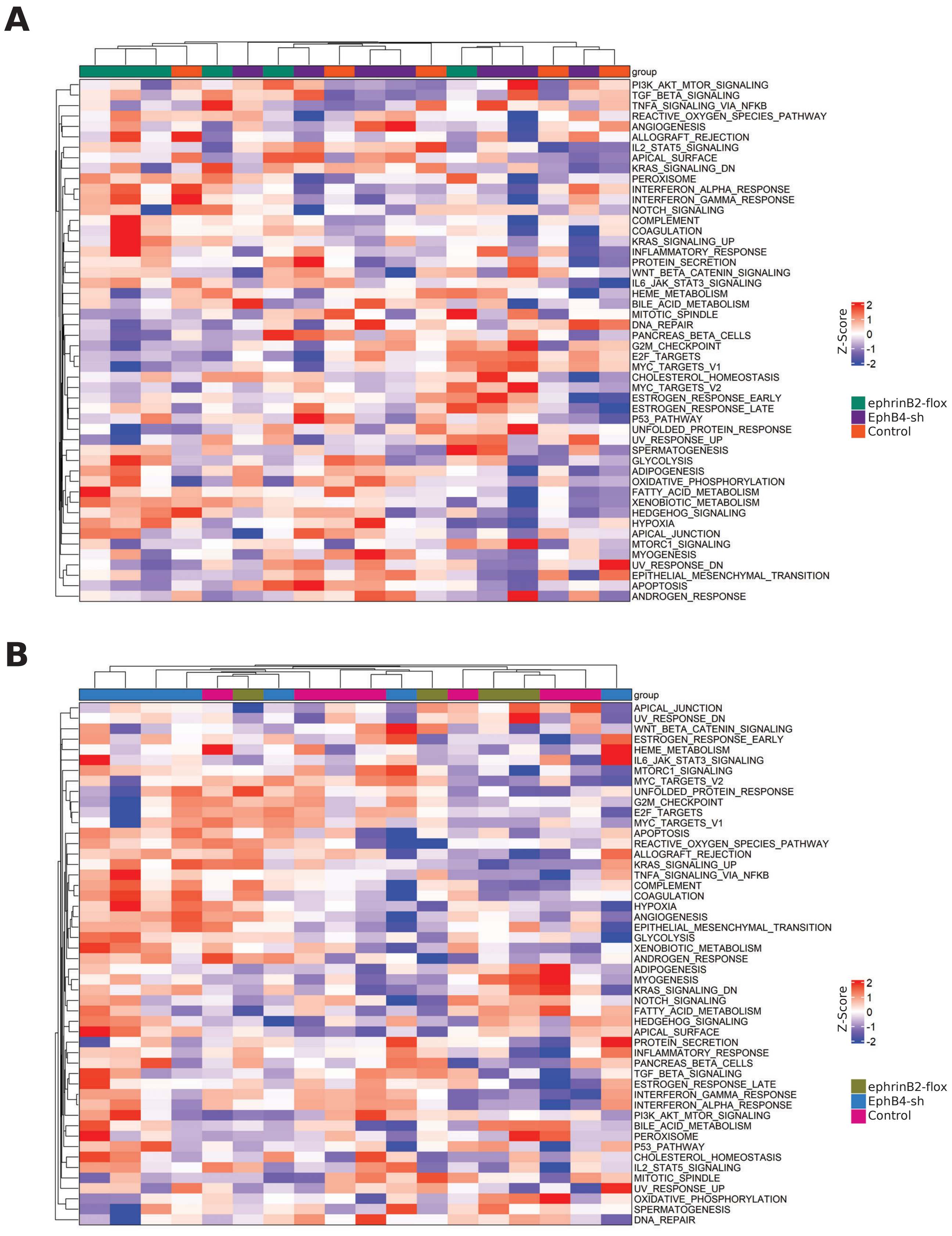


**Supp. Fig. 4: Single-sample GSEA heatmaps.**
(**A**) Heatmap of ssGSEA scores for Hallmark pathways in untreated MOC2 tumors. (**B**) Heatmap of ssGSEA scores for Hallmark pathways in P029 tumors.

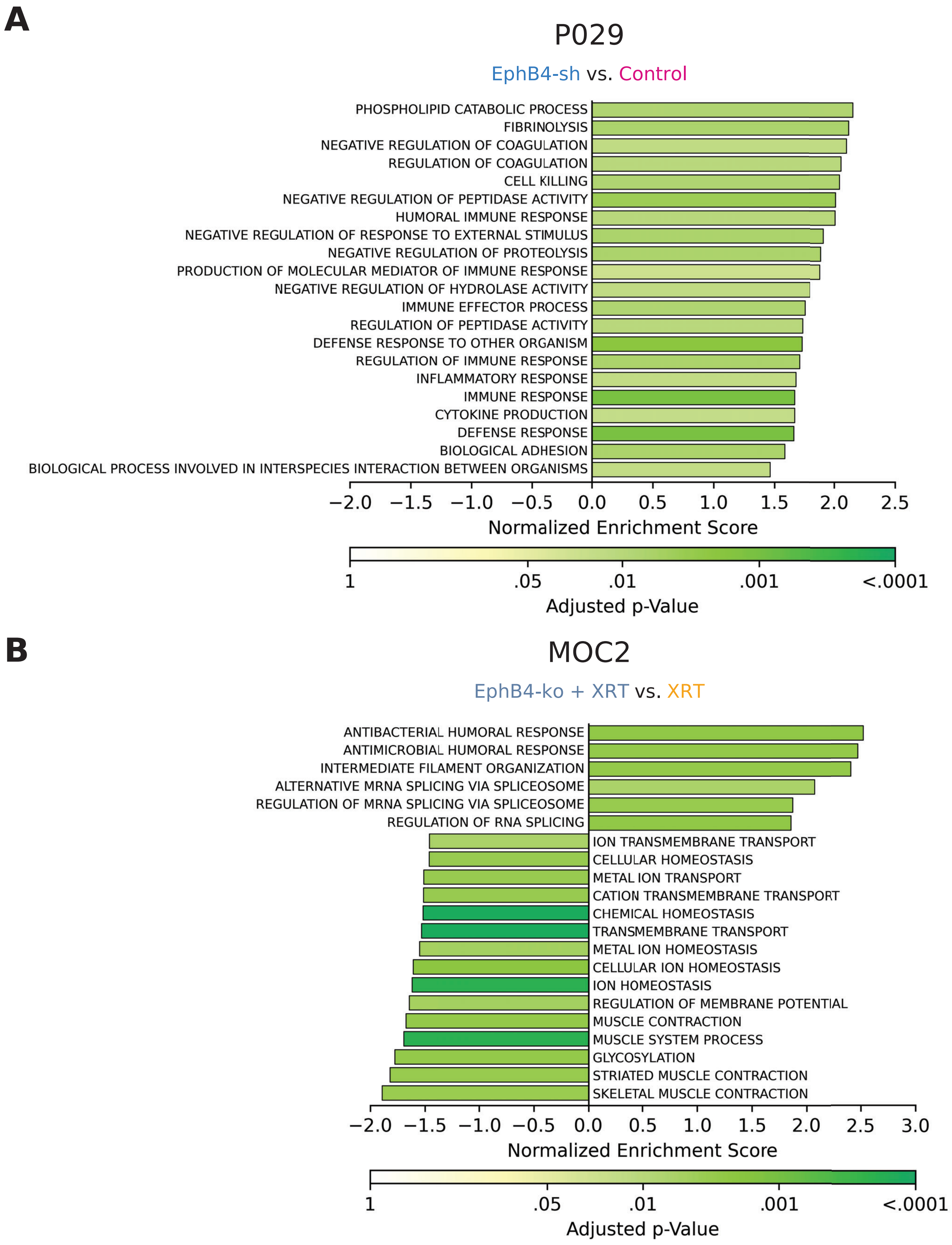


**Supp. Fig. 5: GSEA Results comparing EphB4-knockdown to control MOC2 and P029 tumors.**
(**A**) Normalized enrichment scores from GSEA comparing EphB4 knockdown and control P029 tumors for the top 21 GO Biological Process gene sets by adjusted p-value. (**B**) Normalized enrichment scores from GSEA comparing EphB4 knockdown and control MOC tumors (treated with XRT) for the top 21 GO Biological Process gene sets by adjusted p-value.

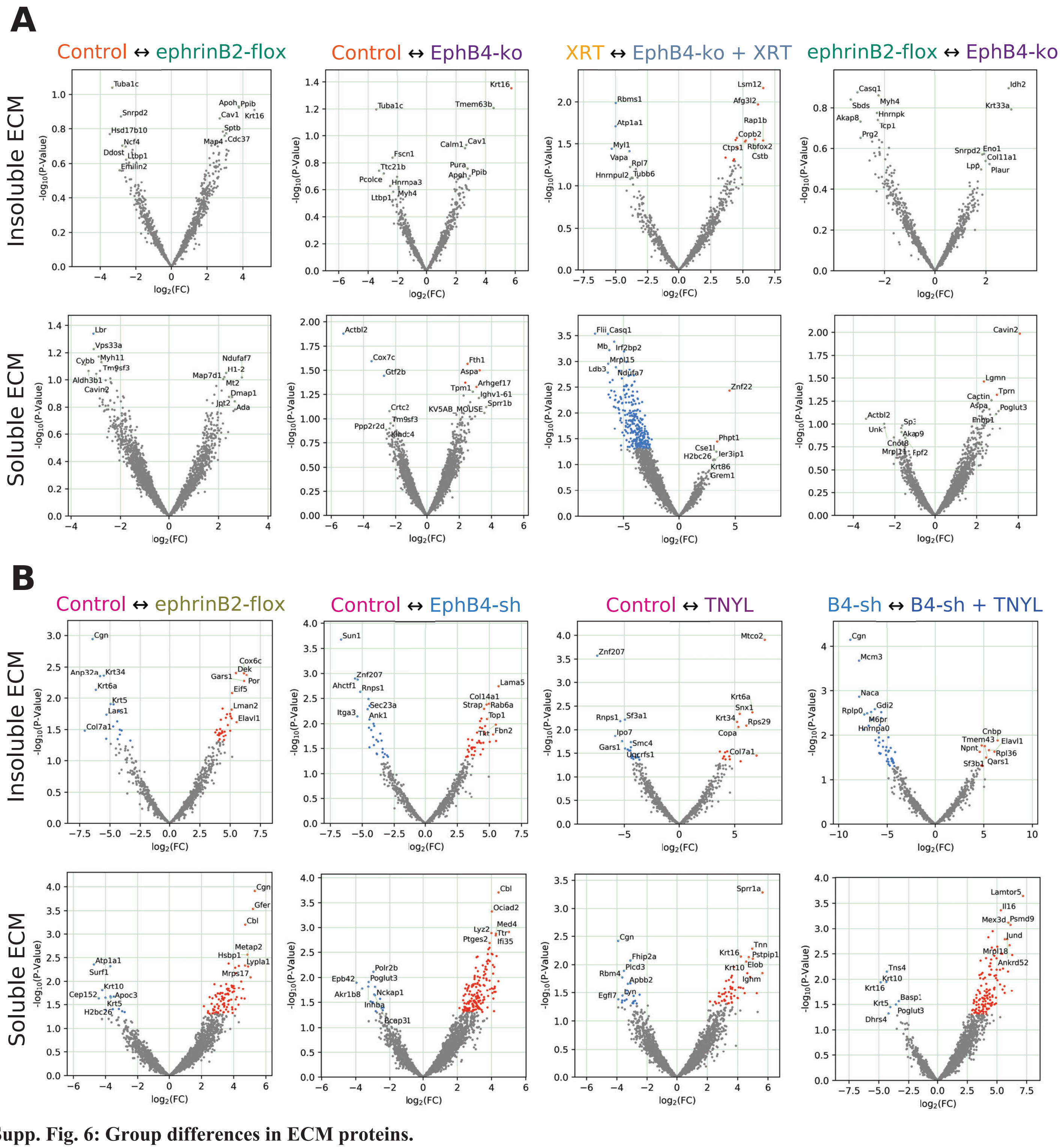


**Supp. Fig. 6: Group differences in ECM proteins.**

(**A**) Volcano plots of differential protein expression for ECM components of MOC2 tumors. The "↔" in each plot title indicates that proteins on the left side of the plot are upregulated in the group to the left and vice versa. (**B**) Volcano plots of differentially protein expression for ECM components of P029 tumors.

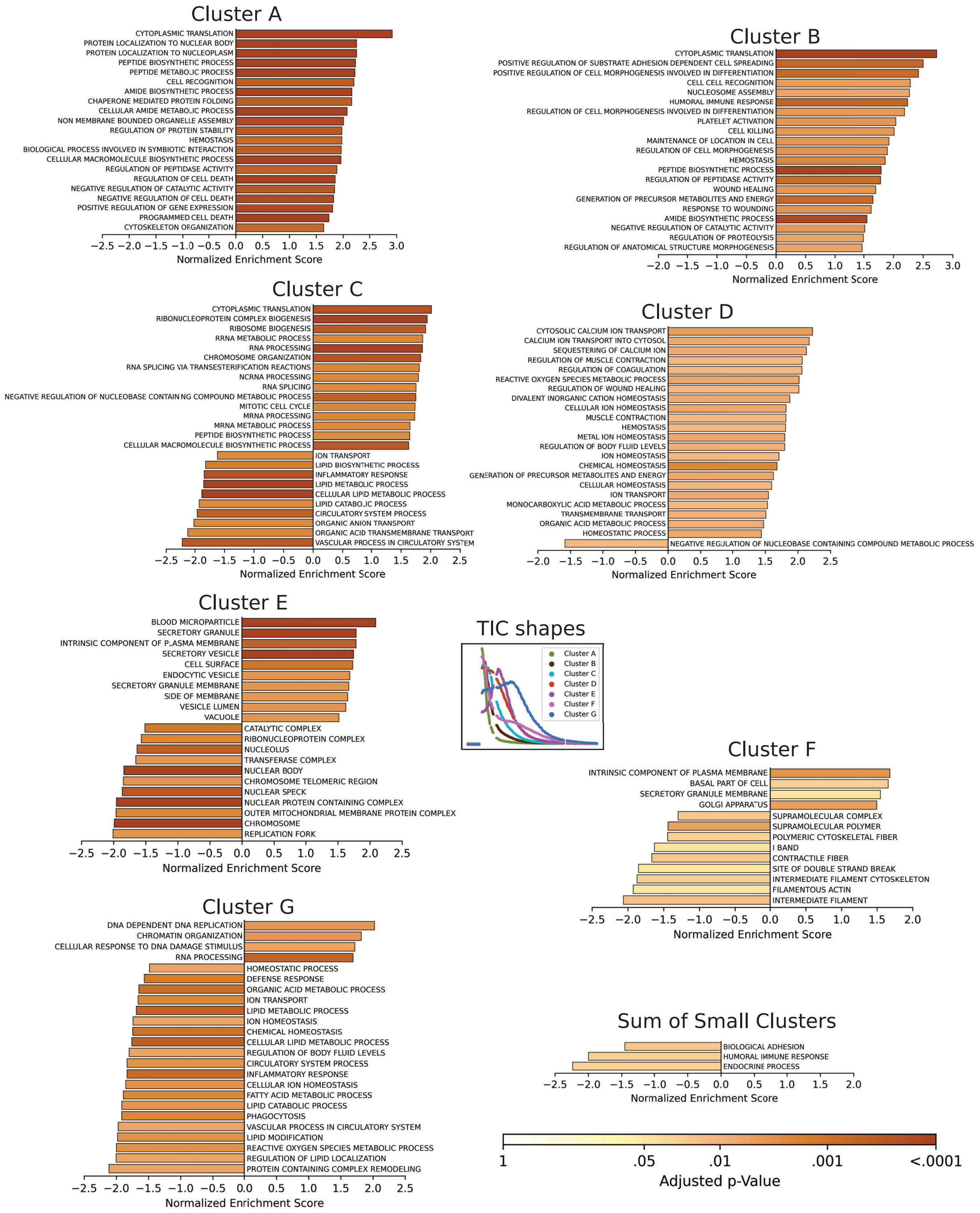


**Supp. Fig. 7: Pathways Correlated with Each TIC Cluster in MOC2 Model.**
Normalized enrichment scores from GSEA (as in **Fig. 5**) for the top 20 GO Biological Process gene sets by adjusted p-value for each named TIC cluster and for the combined area of the remaining TIC clusters (see **Supp. Fig. 2A**). More than 20 may be shown in case of ties or fewer to avoid displaying results with padj > .05. All bars are colored in accordance with the color legend in the lower right. Only MOC2 tumors were included in this analysis.

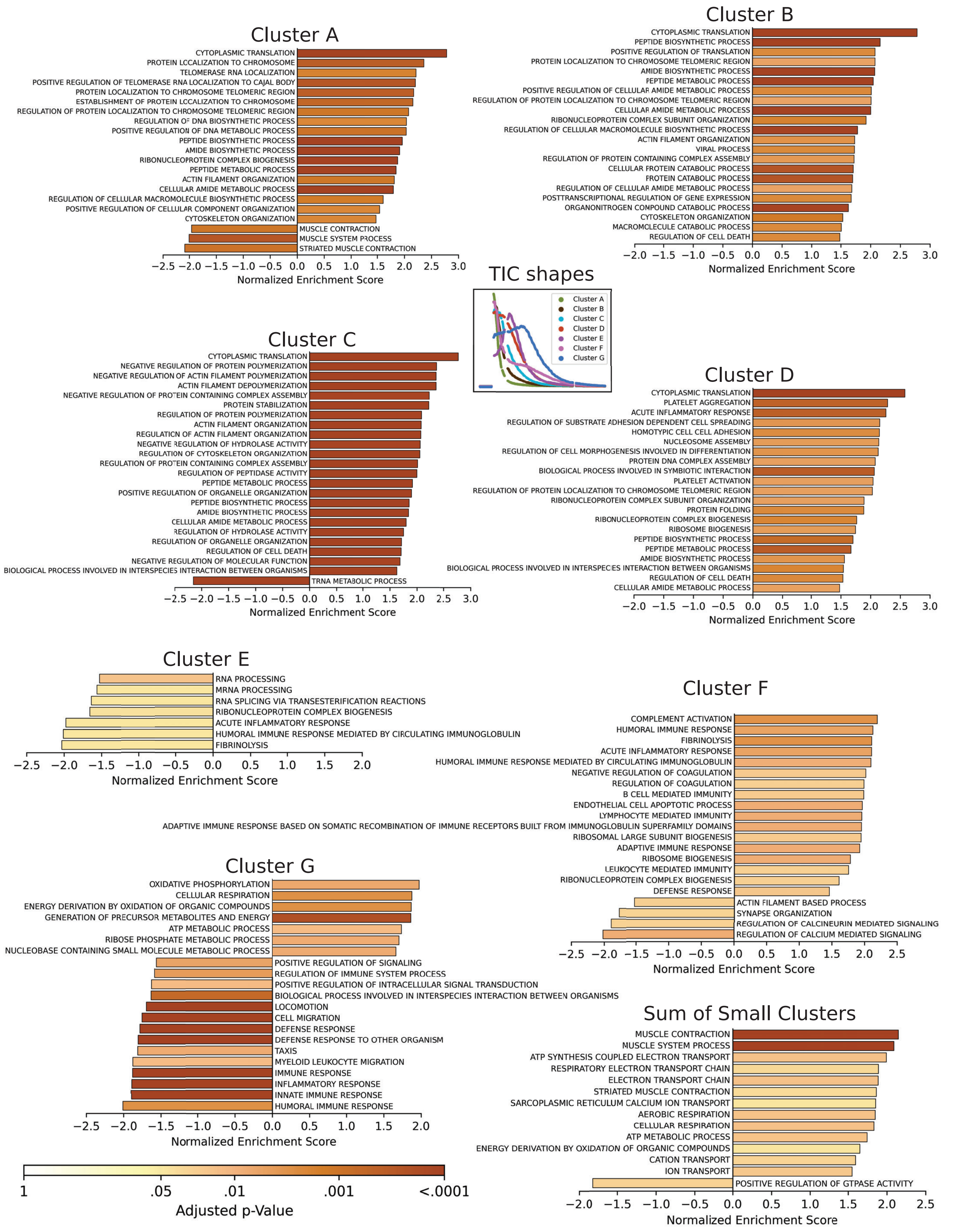


**Supp. Fig. 8: Pathways Correlated with Each TIC Cluster in P029 Model.**
Normalized enrichment scores from GSEA (as in **Fig. 5**) for approximately the top 20 GO Biological Process gene sets by adjusted p-value for each named TIC cluster and for the combined area of the remaining TIC clusters (see **Supp. Fig. 2A**). More than 20 may be shown in case of ties or fewer to avoid displaying results with padj > .05. All bars are colored by adjusted p-value in accordance with the color legend in the lower left. Only P029 tumors were included in this analysis.
